# Transcriptional background effects on a tumor driver gene in a transgenic medaka melanoma model

**DOI:** 10.1101/2022.02.16.480743

**Authors:** Shahad Abdulsahib, William Boswell, Mikki Boswell, Markita Savage, Manfred Schartl, Yuan Lu

## Abstract

The *Xiphophorus* melanoma receptor kinase gene, *xmrk*, is a bona fide oncogene driving melanocyte tumorigenesis of *Xiphophorus* fish. When ectopically expressed in medaka, it not only induces development of several pigment cell tumor types in different strains of medaka, but also induces different tumor types within the same animal, suggesting its oncogenic activity has a transcriptomic background effect. Although the central pathways that *xmrk* utilizes to lead to melanomagenesis are well documented, genes and genetic pathways that modulate the oncogenic effect, and alter the course of disease have not been studied so far. To understand how the genetic networks between different histocytes of *xmrk-*driven tumors are composed, we isolated two types of tumors, melanoma and xanthoerythrophoroma, from the same *xmrk* transgenic medaka individuals, established the transcriptional profiles of both *xmrk-*driven tumors, and compared (1) genes that are co-expressed with *xmrk* in both tumor types, and (2) differentially expressed genes and their associated molecular functions, between the two tumor types. Transcriptomic comparisons between the two tumor types show melanoma and xanthoerythrophoroma are characterized by transcriptional features representing varied functions, indicating distinct molecular interactions between the driving oncogene and the cell type-specific transcriptomes. Melanoma tumors exhibited gene signatures that are relevant to proliferation and invasion while xanthoerythrophoroma tumors are characterized by expression profiles related metabolism and DNA repair. We conclude the transcriptomic backgrounds, exemplified by cell-type specific genes that are downstream of *xmrk* effected signaling pathways, contribute the potential to change the course of tumor development and may affect overall tumor outcomes.

## Introduction

Efforts in the past few decades to identify major disease driver genes have advanced both our understanding of disease etiology and therapeutic development. The genetic background considerably impacts the phenotype of a specific disease. Individuals carrying the same disease driver can exhibit diverged penetrance and expressivity. These effects are linked to genetic background and/or environmental influence on the causal driver function and can complicate diagnosis and proper treatment [1-4]. We now know that genetic background effects are involved in epistatic interactions modulating disease driver function [5, 6]. However, oncogenicity is not universal in different cell types. Known oncogenes preferentially induce certain types of cancer, e.g., *RAS* for pancreas cancer [7], *MYC* for leukemia [8, 9], *SRC* for sarcomas [10], *EGFR* for squamous cell carcinoma, glioblastomas, lung cancer [11-13], *ERBB2* for breast, salivary gland, and ovarian carcinomas [14, 15]. Although the mechanism of cell transformation initiated by the oncogenes are well studied, how they interact with different cell type-specific transcriptomes is not. Delineating interactions between a driving oncogene and a cell-specific transcriptional environment is important for a full understanding of the function, and cell type specific modulators of oncogene action.

Answering the above question, i.e., how oncogenes interact with different transcriptional backgrounds, requires a model system that develops both tractable, and different types of tumors. *Xiphophorus*, a genus of small freshwater fish, is best known for its inter-species hybridization-induced tumorigenesis. It has been shown that a mutant copy of the Epidermal Growth Factor Receptor encoding gene (*egfrb*) named *Xiphophorus* Melanoma Receptor Kinase (*xmrk*) is an oncogene driving tumor development. When this natural mutant gene loses its unlinked regulator following interspecies hybridization due to Mendelian segregation, the *xmrk* overexpresses. It drives tumorigenesis of macromelanophores, a nevus-type of pigment cells in fish. In addition, its level of overexpression correlates with malignancy. Both the histology, and transcriptional features of these pigment cell tumors are similar to human melanoma [16-19]. When the *xmrk* gene is ectopically expressed under a universal promoter in Japanese medaka, a closely related species to *Xiphophorus*, all embryonic cell types underwent dysregulated proliferation and eventually led to embryo death. However, under regulation of the pigment cell-specific *mitfa* promoter, *xmrk* drives several types of pigment cell-specific tumorigenesis. Using the transcriptional signatures that hallmark the *xmrk-*driven tumor, we have developed a platform utilizing gene expression patterns as a phenotype to assess and score anti-cancer drug candidates to perform mid-to high-throughput phenotype drug screening to forward promising chemical lead-structures for further development [20].

Of note, the *xmrk* gene exhibits strong genetic background-dependent tumor phenotypes, as well as diverged tumors from different cell types in the transgenic model: the phenotypes range from xanthoerythrophoromas, extracutaneous melanoma, uveal melanoma in *Carbio* strain; xanthoerythrophoromas, and additional nodular and invasive melanoma in *CabR* strain; extracutaneous melanoma and rarely xanthoerythrophoromas in *HB32C* strain; as well as xanthophore-hyperpigmentation, weakly pigmented melanoma from intestine, and eye melanoma in albino *i-3* strain. The cell types that give rise to these tumors (e.g., dermal and extracutaneous melanocytes for melanoma, xanthophores and erythrophores for xanthoerythrophoromas, uvea pigment cells for eye melanoma) are divergent descendants of neural crest cells. This feature (i.e., different tumor type in the same animal) allows for the identification of shared and diverged gene expression patterns associated with different cell lineages, and to characterize oncogene-transcriptome background interactors (e.g., tumor modifiers) that alter the phenotype of a single driving oncogene.

Therefore, the medaka *xmrk* transgenic model is optimal for studying the question of transcriptional cell-type specific background effect on oncogenes.

Herein, we performed transcriptome profiling of xanthoerythrophoroma and melanoma tumors isolated from the same animals, compared gene expression of the same tumor type among different individuals, and investigated transcriptional differences between tumor types, in order to: 1. Characterize the tumor cell transcriptome to identify the genes that form a network with a single driver oncogene; 2. Investigate transcriptional signatures that differentiate the *xmrk*-driven phenotypes in distinct cell types exhibiting potential diverged transcriptional environments.

## Materials and Methods

### Fish utilized

Seven twelve-month old male *mitf*:*xmrk-*transgenic (*tg-mel*) from the *Carbio* strain were raised and maintained in the *Xiphophorus* Genetic Stock Center in accordance with the Institutional Animal Care and Use Committee (IACUC) protocol (IACUC20173294956). Texas State University has an Animal Welfare Assurance on file (#A4147) with the Office of Laboratory Animal Welfare (OLAW), National Institute of Health.

### RNA isolation

The *tg-mel* medaka were anesthetized by hypothermia, sacrificed, followed by isolation of both melanoma and xanthoerythrophoroma tumors. Tumor samples were immediately placed in 1.5 mL microcentrifuge tubes containing 300 µL TRI Reagent (Sigma Inc., St. Louis, MO, USA) followed by homogenization with a tissue homogenizer. After the initial homogenization, 300 µL of fresh TRI Reagent and 120 µL of chloroform were added to the 1.5 mL microcentrifuge tube and shaken vigorously for 15 sec. Phase separation was performed by centrifugation (12,000 x g for 5 min at 4°C). The aqueous phase was then added to a new 1.5 mL microcentrifuge tube and an additional chloroform extraction was performed (300 µL TRI Reagent, 60 µL chloroform). Following extraction, the nucleic acids were precipitated with 500 µL of 70% EtOH and transferred to a Qiagen RNeasy mini spin column. DNase treatment was performed on-column for 15 min at 25°C, and RNA samples were subsequently eluted with 100 µL RNase-free water. RNA concentrations were quantified with a Qubit 2.0 fluorometer (Life Technologies, Grand Island, NY, USA), and RNA quality was assessed based on RNA integrity (RIN) score with an Agilent 2100 Bioanalyzer (Agilent Technologies, Santa Clara, CA, USA).

### Transcriptional profiling of tumors

All samples sequenced had an RNA Integrity (RIN) score ≥ 8.0. Individual sequencing libraries were constructed using the Illumina TruSeq mRNA Library Prep Kit with polyA selection, and libraries were sequenced (150 bp, paired-end [PE] reads) on the Illumina HiSeq 2000 platform. Raw sequencing reads were subsequently processed using fastx_toolkit for sequencing adaptor removal, low quality base calls, and removal of low-quality sequencing reads,

Processed sequencing reads were mapped to the medaka reference genome (Ensembl release 85, ftp://ftp.ensembl.org/pub/release-85/fasta/oryzias_latipes/dna/) using Tophat2 [21]. Gene expression was quantified using SubReads package function FeatureCounts [22]. Processed read counts per gene are listed in Table S5.

### Principle component analysis and gene co-expression analyses

Gene expression read counts of each sample were normalized to corresponding library size and transcript lengths, and converted to Reads count per Kilobase per Million reads (rpkm). Principle Component Analysis (PCA) was performed using the R package “prcomp” using the scaled rpkm of all samples. Spearman ranking correlation was performed using R programming “cor” function. For each tumor type, samples are ranked based on the *xmrk* expression level. The rpkm values of each gene were subsequently ordered in the ranked samples. A correlation coefficient was subsequently calculated between the gene and *xmrk*. Correlation coefficients > 0.9 or < -0.9 were considered a strong correlation; coefficients between -0.19 and 0.19 were considered no correlation.

### Differentially expressed genes between tumor types

Differentially Expressed Genes (DEGs) between tumor types were identified using R/Bioconductor package edgeR [23]. The log_2_Fold Change (log_2_FC) was calculated using melanoma tumors as control samples. The Area Under Curve (AUC) of the Receiver Operating Characteristic (ROC) curve was calculated to assess true and false positive rates for each gene tested by the R package pROC. A set of statistical thresholds was applied to define DEGs: log_2_FC ≥ 1 or ≤ -1 (log_2_FC ≥ 1 means a gene is higher expressed in xanthoerythrophoromas tumors; log_2_FC ≤ -1 means a gene is higher express in melanoma tumors), log_2_CPM differences ≥ 1, False Discovery Rate (FDR) ≤ 0.05 and ROC curve AUC = 1.

### Functional analyses of genes

Reciprocal Best Hits (RBH) between human orthologs of medaka genes were identified using Blast and subsequently utilized to find human orthologs of medaka genes that co-expressed with *xmrk* or differentially expressed between different tumor types. Gene Set Enrichment Analyses (GSEA) was performed using GSEA tool package in Bioconductor [24, 25]. Medaka datasets were queried against datasets collected in GSEA database. Ingenuity Pathway Analyses (IPA, Qiagen, Redwood City, CA) was used for functional specificity analysis. The goal of these analyses was to identify the biological functions of the *xmrk-*co-expressed genes and inter-tumor type DEGs, therefore, default over-representation analyses was not applied. Genes that were not included and analyzed by either software were manually curated using GeneCard suite [26] and published literature.

## Results

### Tumor type-specific gene co-expression with *xmrk*

Principle Component Analyses (PCA) showed melanoma and xanthoerythrophoromas tumor samples are separated, with the tumor type being the driving dimension that separates the gene expression profiles of all tumor samples (Fig. S1). Genes that are positively or negatively correlated with *xmrk* expression patterns were identified using Pearson correlation in both tumor types. There are 17 genes that are co-expressed with *xmrk* in both tumor types (Fig. 1a; Table S1); 14 genes co-expressed with *xmrk* only in melanoma tumors (Fig. 1b; Table S2); and 29 genes co-expressed with *xmrk* only in xanthoerythrophoroma tumors (Fig. 1c; Table S3). Genes that are co-expressed with *xmrk* in both tumor types are mainly associated with differentiation (*lnx1, lnx2b, pdlim5b* and *sema4b*), proliferation (*dyrk3, egfra, plpp1*), cell cycle regulation (*llgl2*), and cell-microenvironment interaction (*itgb3a*). These genes and related molecular functions represent universal *xmrk* activities regardless of cell types (Fig. 1a; Table S1).

**Figure 1.**
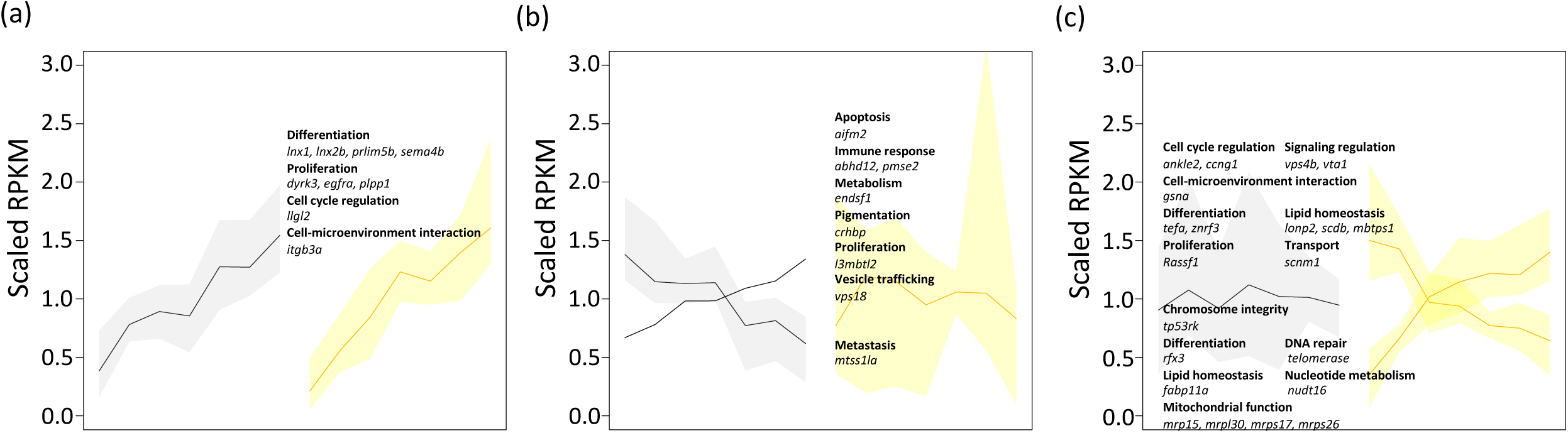
Co-expressed genes of the *xmrk* in melanoma and xanthoerythrophoroma tumors. Genes that exhibit positive or negative correlation to *xmrk* expression in (a) both tumor types, (b) only in melanoma tumors and (c) only in xanthoerythrophoroma tumors are shown. Solid lines represent scaled mean gene expression. Upper and lower boundaries of the shaded areas indicate the max and minimum expression levels respectively.

In contrast, genes that are co-expressed with *xmrk* exclusively in melanoma tumors are associated with apoptosis (*aifm2*), immune response (*abhd12* and *pmse2*), metabolism (*enosf1*), metastasis (*mtss1la*), pigmentation (*crhbp*), proliferation (*l3mbtl2*) and vesicle trafficking [*vps18*; (Fig. 1b; Table S2)]. Genes that co-expressed with *xmrk* only in xanthoerythrophoroma tumors are associated with cell cycle (*ankle2, ccng1*), cell-microenvironment interaction (*gsna*), chromosome integrity (*tp53rk*), differentiation (*rfx3, tefa* and *znrf3*), DNA repair (*telomerase*), fatty acid transportation, metabolism and lipid homeostasis (*lonp2, scdb, fabp11a* and *mbtps1*), mitochondrial function (*mrpl15, mrpl30, mrps17* and *mrps26*), nucleotide metabolism (*nudt16*), proliferation (*rassf1*), transport (*scnm1*), and signaling regulation [*vps4b* and *vta1*; (Fig. 1c; Table S3)].

### Gene expression pattern differentiating tumor types

We assessed differentially expressed genes between melanoma and xanthoerythrophoroma tumors to assess cellular functional differences between the two tumor types. There are 119 genes highly expressed in melanoma tumors, and 63 genes highly expressed in the xanthoerythrophoroma tumors (Fig. 2; Table S4). As expected, genes belonging to pathways associated with eumelanin production are highly expressed in melanoma tumors. We also identified functions of 77 genes that are associated with cell-microenvironment interactions, differentiation, proliferation, metabolism, dopamine homeostasis, immune response and PPAR/RXR activation. All proliferation related genes, and a majority of genes associated with differentiation, and cell-microenvironment related are higher expressed in melanoma tumors than xanthoerytrhophoromas (Fig. 3).

**Figure 2.**
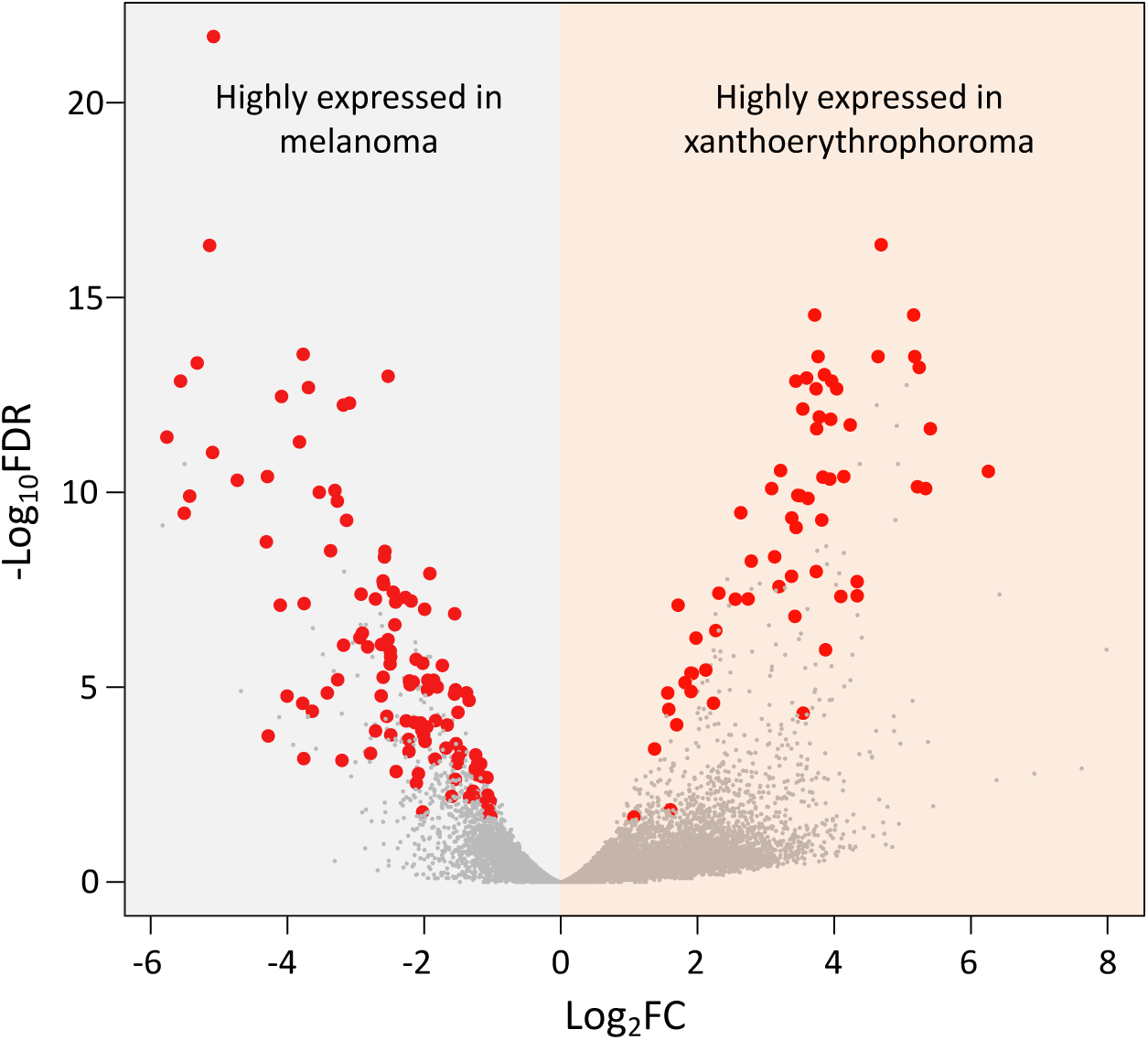
Differentially expressed genes between melanoma and xanthoerythrophoroma tumors. Differentially expressed genes are identified between melanoma and xanthoerythrophoroma tumors. There are 119 genes highly expressed in the melanoma tumors, and 63 genes highly expressed in the xanthoerythrophoroma tumors. Volcano plot shows log_2_FC between tumor types, and -log_10_FDR of differential expression test. Red dots highlight differentially expressed genes, gray dots are genes that are not differentially expressed.

**Figure 3.**
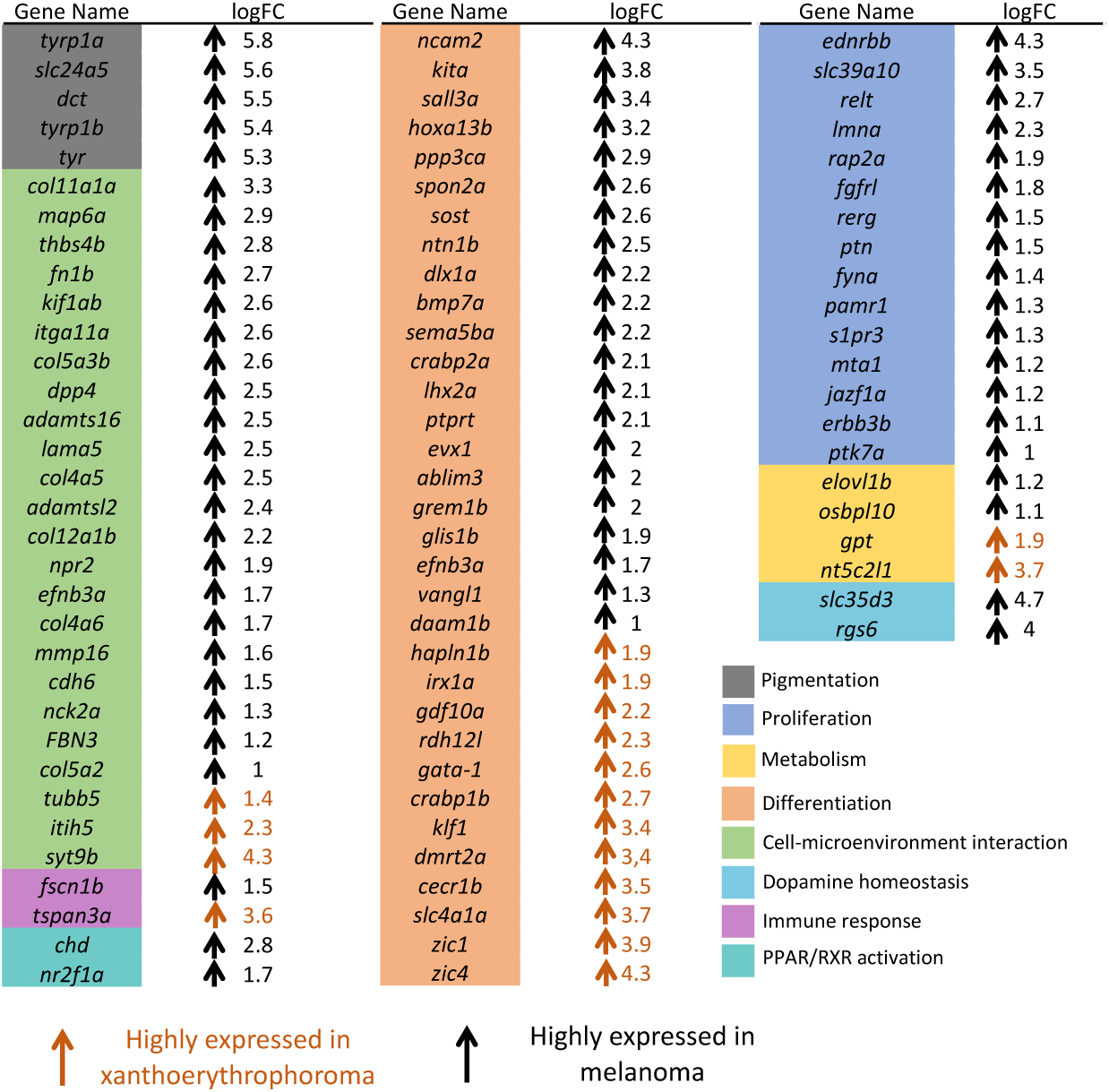
Functional categories of differentially expressed genes between tumor types. Functions of inter-tumor type differentially expressed genes are shown. Colored blocks represent functional categories. Black arrows mean a specific gene over expressed in melanoma, and orange arrows mean a gene over expressed in xanthoerythrophoroma tumors, with the numbers indicated Log_2_FC of the relative gene expression.

## Discussion

The *xmrk* is a bona fide oncogene. It is a duplicated mutant *egfr* copy in a few species belonging to the Central American fish genus *Xiphophorus*, and it drives spontaneous tumorigenesis in interspecies *Xiphophorus* hybrids due to negative epistasis. Functional studies on *xmrk* ectopically expressed in-vitro in murine cells or in transgenic zebrafish and medaka, revealed it drives dedifferentiation, enhanced proliferation, and tumorigenesis [27-30]. Combined with tumor transcriptome, the tumor phenotypical differences allow for the deconvolution of transcriptional networks that interact with *xmrk* to modify its function. The *xmrk* is constitutively active independent of EGF binding, due to mutation induced dimerization [31]. Since human EGFR is associated to a majority types of human cancers, characterizing its modifiers is important in fully understanding its mode of action, and overcoming current therapeutic resistance to anti-EGFR compounds.

The recurrent somatic mutations in tumor cells affect almost every level of transcriptional control (e.g., cellular signaling pathways, transcription factors, enhancer, chromatin structure) [32-39]. Oncogenes can disrupt normal gene expression regulatory mechanisms and transform normal cells into cancer cells. Even though tumorigenesis is a multi-step process, oncogene expression can be indispensable for cancer cell proliferation even the cancer cells have progressed long after a neoplastic state. This proliferative reliance on oncogene expression is named oncogene transcriptional addiction [32, 40]. Studies of *xmrk-*driven tumors in both *Xiphophorus* and transgenic medaka showed tumor cells exhibit high levels of *xmrk* expression [29, 41]. The *xmrk-*driven cancer can be observed 4 weeks following hatching in *xmrk-*medaka. The fish utilized in this study are one-year old with advanced stage tumors, suggesting the tumor cells are addicted to *xmrk* expression, and the transcriptome of the cancer cells is still directly under master regulation of *xmrk* despite exhaustion of *xmrk’*s initial neoplasm triggering activity.

The medaka transgenic system enables investigation of how *xmrk* interacts with transcriptomes of different cell lineages, and characterize cell type-specific genes interactions. Using the *xmrk-*transgenic medaka system, we sought to answer two questions: 1. How are the genetic networks under *xmrk* regulation different between the two tumor types that are driven by the same oncogene; 2. How are the transcriptional phenotypes different between tumor types as a result of driving oncogene *xmrk* interaction with the cell type specific transcriptional landscape.

To answer the first question, we compared genes that co-expressed with *xmrk* in both melanoma and xanthoerythrophoroma tumors. Although *xmrk* drives proliferation, cell-microenvironment interaction, cell cycle and differentiation in both tumor types, the observation of varied genetic functions that are associated with genes that co-expressed with *xmrk* in a tumor type-specific way is suggestive that *xmrk* regulate different cellular processes between melanoma and xanthoerythrophoroma tumor cells.

To answer the second question, we compared bulk transcriptomic differences between the melanoma and xanthoerythrophoroma tumors. In consistence with the distinguished coloration between the two tumor types, genes associated with pigmentation pathways are observed. Differentiation related genes are also a reflection of cell type difference between the two tumor types. However, presence of pivotal differentiation genes (33 genes; Fig. 3) suggest the two tumor types may be at varied differentiation stages or potential. Genes related to proliferation and cell-microenvironment interactions are predominantly highly expressed in melanoma tumors. These include a few proto-oncogenes like endothelin receptor *ednrbb*, sarcoma viral oncogene homolog *kita*, Ras like estrogen regulated growth factor *rerg*, pleiotrophin *ptn*, FYN proto-oncogene *fyna*, erbb2 receptor tyrosine kinase 3b *erbb3b*. This evidence, along with previous observation that melanotic tumors are highly invasive into musculature and internal organs while xanthoerythrophoromas grow more as epiphytic nodules [42], suggests the melanoma tumors are more proliferative than xanthoerythrophoromas. Some of these proto-oncogenes are known to be induced by *xmrk* in *Xiphophorus* [43-46], culture murine cells, and most of them were reported to be involved in human melanomagenesis [43, 47-52]. For example, the *fyna* mouse ortholog (Fyn) has been shown to play an important role in *xmrk* signal transduction at protein and post-translational level [19]. Herein this study we confirm the activity is also displayed at transcriptional level. It is also important to note that fibronectins *fn1b*, integrin *itga11a*, laminin *lama4*, cadherin *cdh6*, collagens *col11a1a, col5a3b, col4a5, col12a1b, col4a6 col5a2*, matrix metalloproteinase *mmp16*, ADAMs *adamts16* and *adamtsl2* are also highly expressed in the melanoma tumors. These genes are reliable markers for tumor cell invasion and metastasis. For example, *col4a5* encodes collagen that make up basement membrane that are involved in metastasis [53]; *cdh6* involves in epithelial-mesenchymal transition [54, 55]; *mmp16* promotes tumor metastasis [56]. Combining this observation with the highly expressed proto-oncogenes suggests the melanoma tumors exhibit a higher potential to invasion.

In summary, genes that are co-expressed with *xmrk* in melanoma and xanthoerythrophoroma tumors, and differentially expressed genes between the two tumor types are involved in diverged biological functions as a result of distinct molecular interactions between the driving oncogene and cell type specific modifiers. We conclude *xmrk* oncogene exhibits a strong transcriptomic background dependent activity. The oncogene modifiers can change the course of tumor development and may affect overall tumor outcomes.

## Acknowledgements

This work was supported by National Institutes of Health grant R24-OD-011120, and R15-CA-223964.

## Figure Legend

**Supplement Table S1. Genes exhibiting positive or negative correlation to**

***xmrk* expression in both tumor types**

**Supplement Table S2. Genes exhibiting positive or negative correlation to**

***xmrk* expression only in melanoma tumors**

**Supplement Table S3. Genes exhibiting positive or negative correlation to**

***xmrk* expression only in xanthoerythrophoroma tumors**

**Supplement Table S4. Differentially expressed genes between melanoma and xanthoerythrophoroma tumors**

**Supplement Table S5. RPKM values of gene expression Supplement Figure**

**Figure S1. Principle component analyses of gene expression profiles** Scatter plot showing distribution of samples distribution on principle component (PC) 1 & 2. Black square dots represent melanoma tumors, and orange square dots represent xanthoerythrophoroma tumors.

## Additional Information

The authors have no competing interests, or other interests that might be perceived to influence the results and/or discussion reported in this paper.

## Data availability statement

The raw sequencing files are deposited in Gene Expression Omnibus. The accession number will be publicly available upon manuscript acceptation for publication.

